# Growth pattern Learning for Unsupervised Extraction of Cancer Kinetics

**DOI:** 10.1101/2020.06.13.140715

**Authors:** Cristian Axenie, Daria Kurz

**Affiliations:** *Audi Konfuzius-Institut Ingolstadt Lab*, Technische Hochschule Ingolstadt, Ingolstadt, Germany; *Interdisciplinary Breast Center*, Helios Clinic Munich West, Munich, Germany

**Keywords:** Unsupervised Learning, Neural Networks, Cancer Kinetics, Tumor Growth

## Abstract

Neoplastic processes are described by complex and heterogeneous dynamics. The interaction of neoplastic cells with their environment describes tumor growth and is critical for the initiation of cancer invasion. Despite the large spectrum of tumor growth models, there is no clear guidance on how to choose the most appropriate model for a particular cancer and how this will impact its subsequent use in therapy planning. Such models need parametrization that is dependent on tumor biology and hardly generalize to other tumor types and their variability. Moreover, the datasets are small in size due to the limited or expensive measurement methods. Alleviating the limitations that incomplete biological descriptions, the diversity of tumor types, and the small size of the data bring to mechanistic models, we introduce Growth pattern Learning for Unsupervised Extraction of Cancer Kinetics (GLUECK) a novel, data-driven model based on a neural network capable of unsupervised learning of cancer growth curves. Employing mechanisms of competition, cooperation, and correlation in neural networks, GLUECK learns the temporal evolution of the input data along with the underlying distribution of the input space. We demonstrate the superior accuracy of GLUECK, against four typically used tumor growth models, in extracting growth curves from a four clinical tumor datasets. Our experiments show that, without any modification, GLUECK can learn the underlying growth curves being versatile between and within tumor types.

## 1 Background

Ideally, if we would have understood all tumor biology from neoplastic cells to metastatic cancer, we could build a model that would anticipate the development curve of a tumor into what’s to come given its present state. Of course, we cannot do that because our knowledge is sadly incomplete. Contributing to this is the fact that repeated measures of growing tumors - required for any detailed study of cancer kinetics - are very difficult to obtain.

Tumor growth can typically be described in three frameworks: 1) in vitro [14]; 2) exploratory in vivo frameworks (animal models with implanted tumors, [26,6]); or 3) clinical tumor growth using non intrusive imaging strategies, [9]. But, even in the best scenarios, obtaining accurate size measures of the irregular, three-dimensional masses without influencing the physiology of either tumor or host remains problematic. Nevertheless, such an assessment is crucial as it can subsequently guide cancer therapy.

Mostly due to such limitations, theorists sought “growth laws” that seek to represent general tumor kinetics without reference to any particular tumor type.

### 1.1 Tumor growth and its implications

Extensive biological studies have been devoted to tumor growth kinetics, addressing both mass and volume evolution. Such studies postulated that, principles of tumor growth might result from general growth laws, typically amenable to differential equations [12]. These models have a twofold objective: 1) testing growth hypotheses or theories by assessing their descriptive power against experimental data and 2) estimating the prior or future course of tumor progression. Such an assessment can be used as a personalized clinical prediction tool [3], or in order to determine the efficacy of a therapy [7]. With GLUECK we try to address the two objectives by exploiting the benefits of learning and generalization in neural networks.

### 1.2 Mechanistic models of tumor growth

Different tumor growth patterns have been identified experimentally and clinically and modelled over the years. A large number of ordinary differential equations (ODE) tumor growth models [12] have been proposed and used to make predictions about the efficacy of cancer treatments.

The first theory that formulated a consistent growth law implied that tumors grow exponentially, a concept that obviously stems from the “uncontrolled proliferation” view of cancer. However, this hypothesis is wrong. Assuming that a tumor increases exponentially, then a one-dimensional estimate of its size (e.g. diameter) will also increase. In other words, both volume and “diameter” log plots over time are linear, if and only if the tumor is exponentially increasing. But, detailed studies of tumor growth in animal models [23] reveal that the tumor “diameter” appears to be linear, not its logarithm, suggesting a decreasing volumetric growth rate over time, Figure 1. There probably is no single answer to the question, which is the best model to fit cancer data [17]. In our analysis we chose four of the most representative and typically used scalar growth models, namely Logistic, von Bertalanffy, Gompertz and Holling, described in Table 1 and depicted in Figure 2. Despite their ubiquitous use, in either original form or embedded in more complex models [18], the aforementioned scalar tumor growth models are confined due to: a) the requirement of a precise biological description (i.e. values for *α,β*, λ and *k* correspond to biophysical processes); b) incapacity to describe the diversity of tumor types (i.e. each is priming on a type of tumor), and c) the small amount and irregular sampling of the data (e.g. 15 measurements with at days 6, 10, 13, 17, 19, 21, 24, 28, 31, 34, 38, 41, 45, 48 for breast cancer growth in [26]).

**Fig.1.**
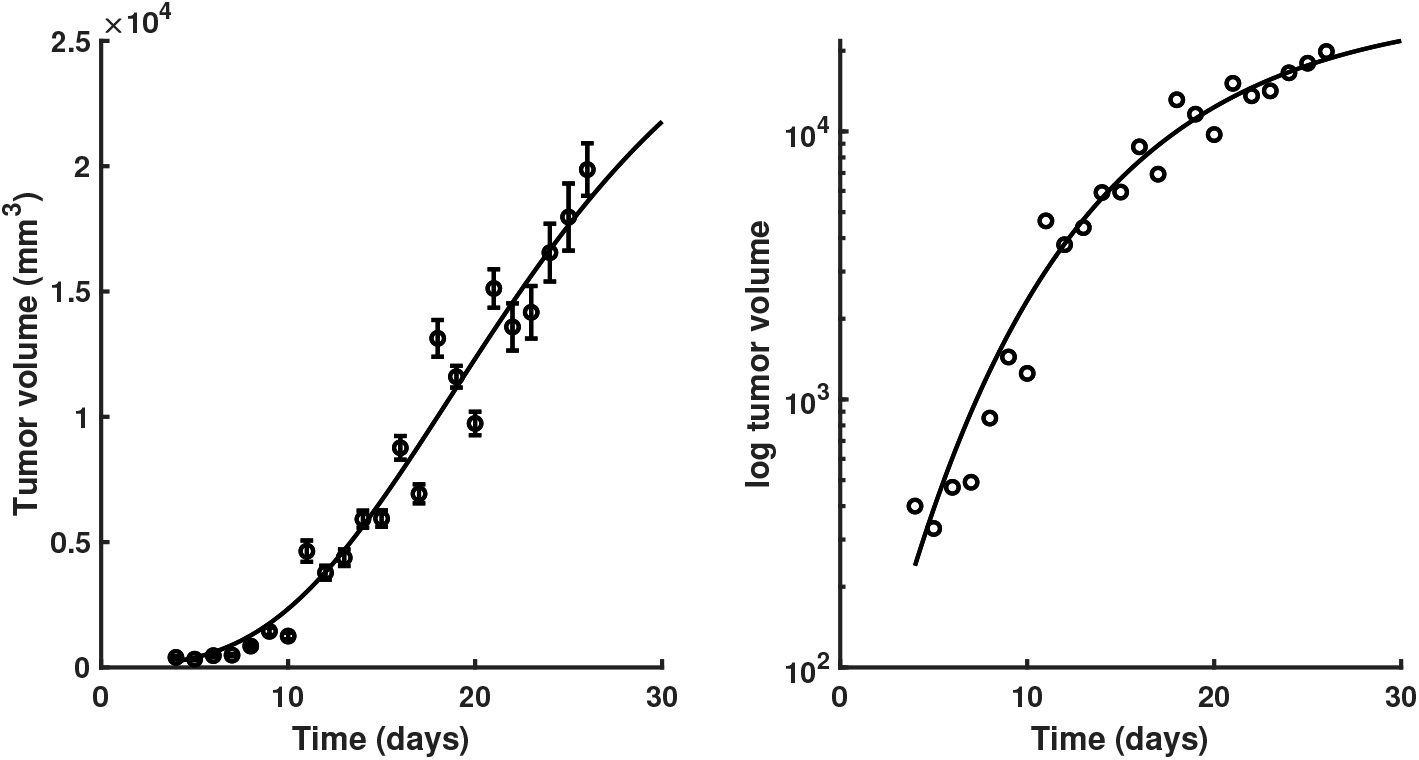
Growth kinetics of Fortner Plasmacytoma 1 tumors. Points represent mean volume of subcutaneous tumor implants in mice, error bars represent +/-1 standard error of the mean at each point. Data from [23]

**Table 1.** Overview of tumor growth models in our study. Parameters: *N* - cell population size (or volume / mass thorough conversion [15]), *α* - growth rate, *β* - cell death rate, λ - nutrient limited proliferation rate, *k* - carrying capacity of cells.

| Model | Equation |
| --- | --- |
| Logistic [25] | $\frac{dN}{dt} = \alpha N - \beta N^2$ |
| Bertalanffy [27] | $\frac{dN}{dt} = \alpha N^\lambda - \beta N$ |
| Gompertz [13] | $\frac{dN}{dt} = N(\beta - \alpha \ln N)$ |
| Holling [2] | $\frac{dN}{dt} = \frac{\alpha N}{k+N} - \beta N$ |

**Fig.2.**
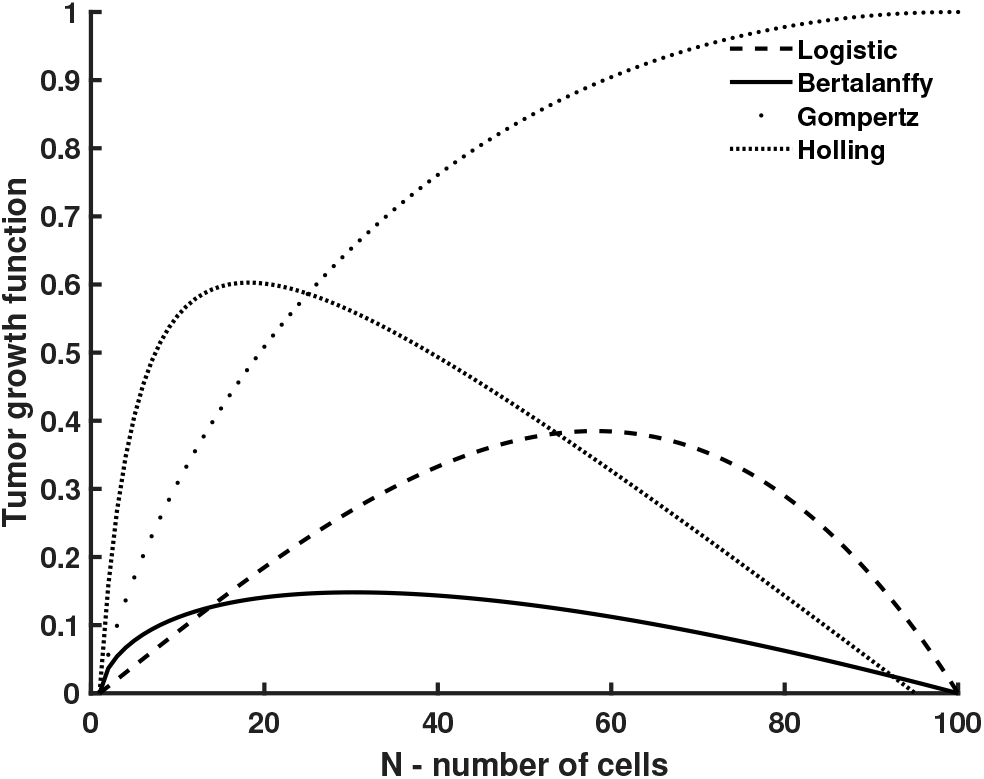
Comparison of four tumor growth rate functions.

Although such competing canonical tumor growth models are commonly used, how to decide which of the models to use for which tumor types is still an open question. In an attempt to answer this questions, the work in [22] built a broad catalogue of growth laws and recommendations on the best fit and parameter ranges. Now, assuming that a set of models was selected for a treatment, the next question is how to quantify their differences in predictions, for instance, in the presence and absence of chemotherapy? Addressing this final question, the study in [19] showed that tumor growth model selection can lead to as much as twelve-fold change in estimated outcomes, and that the model that best fits experimental data might not be the best model for future growth. To cope with these inherent limitations, predictive models of tumor growth were developed.

### 1.3 Predictive models of tumor growth

Several researchers have tried to find the “right” ODE growth law by fitting different models onto a limited number of experimental tumor growth data sets [22]. All in all, the findings are somewhat inconclusive, with findings indicating that the choice of growth model is at least partially dependent on the type of tumor [5]. Addressing this challenge multiple studies “augmented” traditional growth models with predictive capabilities wrapping their parameter estimation into probabilistic or machine learning frameworks. We comparatively review three most relevant approaches.

Building upon the basic growth models (i.e. adapted Gompertz model on tumor phases) and vascular endothelial growth factor (VEGF), the work in [11] trained and validated a model using published in vivo measurements of xenograft tumor volume. The model employed Nonlinear Least Squares (NLS) optimization together with a global sensitivity analysis to determine which of the four tumor growth kinetic parameters impacts the predicted tumor volume most significantly. The major disadvantage of the method is that NLS requires iterative optimization to compute the parameter estimates. Initial conditions must be reasonably close to the unknown parameter estimates or the optimization procedure may not converge. Moreover, NLS has strong sensitivity to outliers. GLUECK alleviates the need of expensive optimization through a learning mechanism on multiple timescales, that is guaranteed to converge [4]. Instead of selecting best model parameters that describe a certain dataset, GLUECK uses a data-driven learning and adaptation to the data distribution (see Methods) implicitly achieving this.

Using growth simulations on functional Magnetic Resonance Imaging (fMRI) and Partial Differential Equations (PDE)-constrained optimization, the work in [1] used discrete adjoint and tangent linear models to identify the set of parameters that minimize an objective functional of tumor cell density and tissue displacement at a specific observation time point. Conversely, GLUECK does not depend on expensive imaging data and a precise PDE modelling of the problem, rather it exploits the data to extract the underlying growth pattern through simple and efficient operations in neural networks (see Methods).

Finally, the work in [24] performed a quantitative analysis of tumor growth kinetics using nonlinear mixed-effects modeling on traditional mechanistic models. Using Bayesian inference, they inferred tumor volume from few (caliper and fluorescence) measurements. The authors proved the superior predictive power of the Gompertz model when combined with Bayesian estimation. Such work motivated the need for GLUECK. This is mainly because GLUECK: 1) departs completely from the limitations of a mechanistic model of tumor growth (e.g. Gompertz) and 2) it alleviates the difficulty of choosing the priors, the computational cost at scale, and the incorporation of the posterior in the actual prediction - that typically affect Bayesian methods.

### 1.4 Peculiarities of tumor growth data

Before we introduce GLUECK we strengthen our motivation for this work by iterating through the peculiarities of tumor growth data.

Tumor growth data:

is **small**, only a few data points with, typically, days level granularity, [21].
is **unevenly sampled**, with irregular spacing among measurements [26].
data has **high variability** between and within tumor types (e.g. breast versus lung cancer, [5] and each type of treatment (i.e. administered drug) modifies the growth curve [11].
is **heterogeneous** and sometimes **expensive to obtain** (e.g.volume assessed from bio-markers, fMRI [1], fluorescence imaging [20], flow cytometry, or calipers [6]).
poses **challenges in selecting the best model** [1,11,19].
**determines cancer treatment planning** [22].

## 2 Materials and methods

In the current section we describe the underlying functionality of GLUECK along with the datasets used in our experiments.

### 2.1 Introducing GLUECK

GLUECK is an unsupervised learning system based on Self-Organizing Maps (SOM) [16] and Hebbian Learning (HL) [8] as main ingredients for extracting underlying relations among correlated timeseries. In order to introduce GLUECK, we provide a simple example in Figure 3. Here, we consider data from a cubic growth law (3^rd^ powerlaw) describing the impact of sequential dose density over a 150 weeks horizon in adjuvant chemotherapy of breast cancer [10]. In this simple example, the two input timeseries (i.e. the cancer cell number and the irregular measurement index over the weeks) follow a cubic dependency (cmp. Figure 3a). When presented the data, GLUECK has no prior information about timeseries’ distributions and their generating processes, but learns the underlying (i.e. hidden) relation directly from the input data in an unsupervised manner.

**Fig.3.**
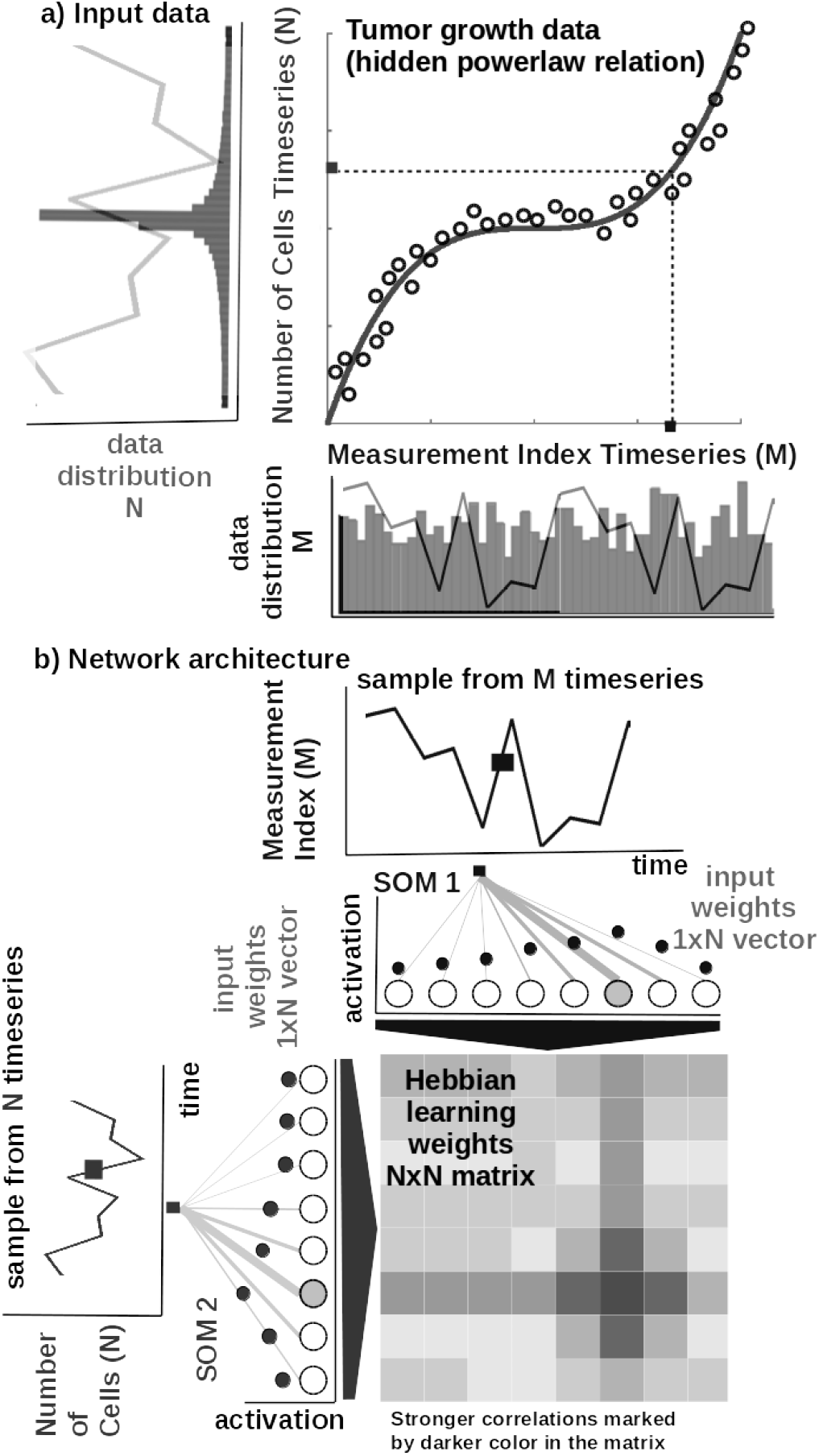
Basic functionality of GLUECK. a) Tumor growth data resembling a non-linear relation and its distribution - relation is hidden in the timeseries (i.e. number of cells vs. measurement index). Data from [10]. b) Basic architecture of GLUECK: 1D SOM networks with *N* neurons encoding the timeseries (i.e. number of cells vs. measurement index), and a *NxN* Hebbian connection matrix coupling the two 1D SOMs that will eventually encode the relation.

#### Core model

The input SOMs (i.e. 1D lattice networks with *N* neurons) are responsible to extract the distribution of the incoming sensory data, depicted in Figure 3a, and encode sensory samples in a distributed activity pattern, as shown in Figure 3b. This activity pattern is generated such that the closest preferred value of a neuron to the input sample will be strongly activated and will decay, proportional with distance, for neighbouring units. This accounts for the basic quantization capability of SOM and augments it with a dimension corresponding to latent representation resource allocation (i.e. number of neurons allocated to represent the input space). The SOM specialises to represent a certain (preferred) value in the sensory space and learns its sensitivity, by updating its tuning curves shape. Given an input sample *s^p^*(*k*) from one timeseries at time step *k*, the network follows the processing stages depicted in Figure 4. For each *i*-th neuron in the *p*-th input SOM, with preferred value 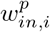 and tuning curve size 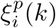, the elicited neural activation is given by

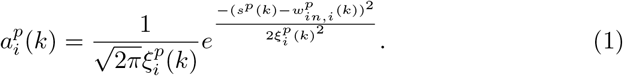

The winning neuron of the *p*-th population, *b^p^*(*k*), is the one which elicits the highest activation given the timeseries sample at time *k*

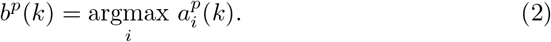

The competition for highest activation in the SOM is followed by cooperation in representing the input space. Hence, given the winning neuron, *b^p^*(*k*), the cooperation kernel,

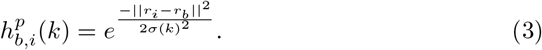

allows neighbouring neurons (i.e. found at position *r_i_* in the network) to precisely represent the input sample given their location in the neighbourhood *σ*(*k*) of the winning neuron. The neighbourhood width *σ*(*k*) decays in time, to avoid twisting effects in the SOM. The cooperation kernel in Equation 3, ensures that specific neurons in the network specialise on different areas in the input space, such that the input weights (i.e. preferred values) of the neurons are pulled closer to the input sample,

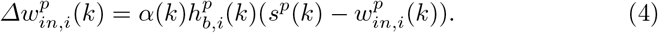

**Fig.4.**
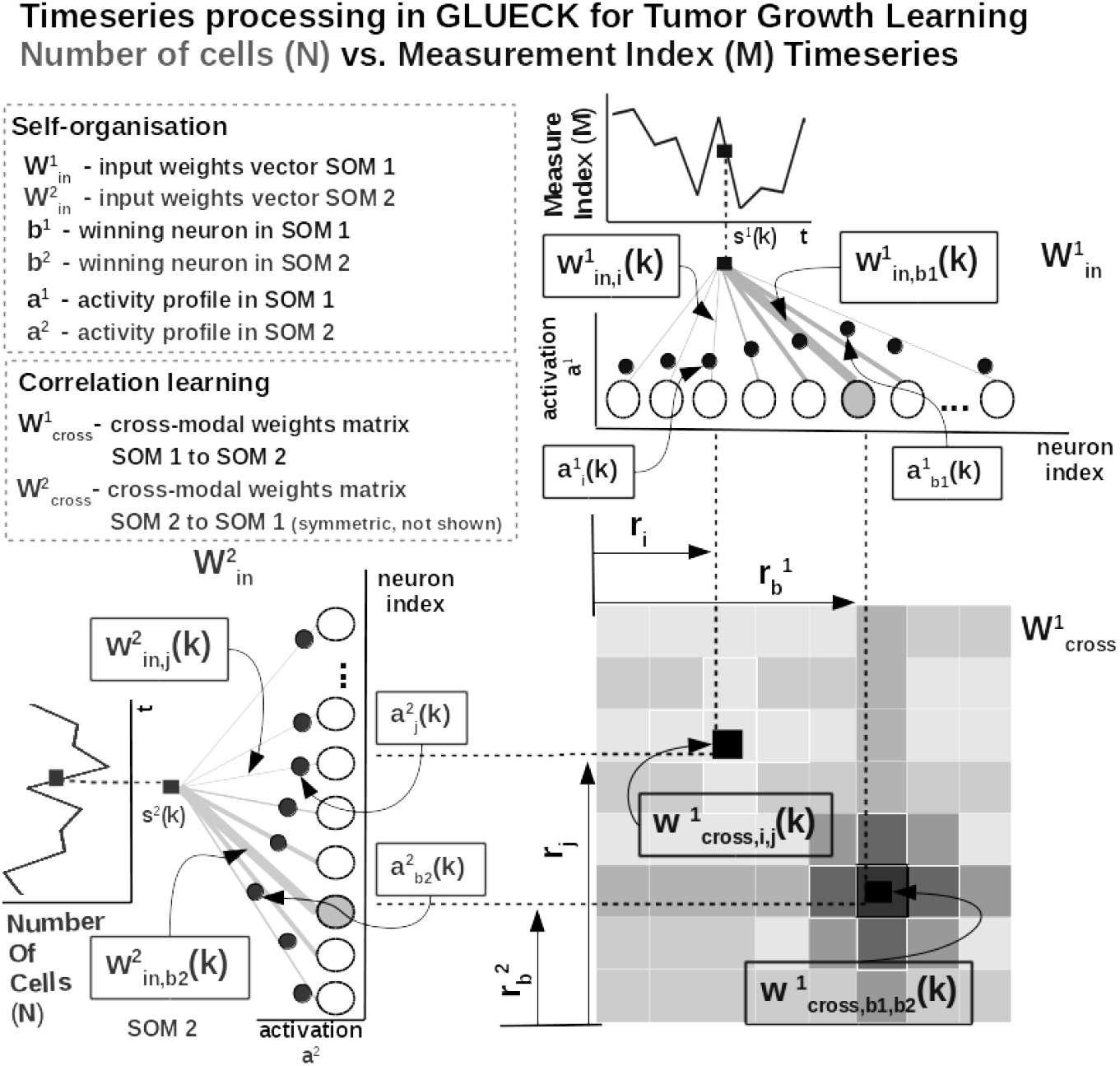
Detailed network functionality of GLUECK, instantiated for tumor number of cells vs. measurement index data from [10].

This corresponds to updating the tuning curves width 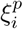 as modulated by the spatial location of the neuron in the network, the distance to the input sample, the cooperation kernel size, and a decaying learning rate *α*(*k*),

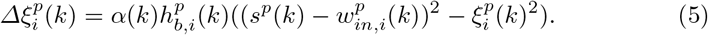

As an illustration of the process, let’s consider learned tuning curves shapes for 5 neurons in the input SOMs (i.e. neurons 1, 6, 13, 40, 45) encoding the breast cancer cubic tumor growth law, depicted in Figure 5. We observe that higher input probability distributions are represented by dense and sharp tuning curves (e.g. neuron 1, 6, 13 in SOM1), whereas lower or uniform probability distributions are represented by more sparse and wide tuning curves (e.g. neuron 40, 45 in SOM1). Using this mechanism, the network optimally allocates resources (i.e. neurons). A higher amount of neurons to areas in the input space, which need a finer resolution; and a lower amount for more coarsely represented areas. Neurons in the two SOMs are then linked by a fully (all-to-all) connected matrix of synaptic connections, where the weights in the matrix are computed using Hebbian learning. The connections between uncorrelated (or weakly correlated) neurons in each population (i.e. *w_cross_*) are suppressed (i.e. darker color) while correlated neurons connections are enhanced (i.e. brighter color), as depicted in Figure 4. Formally, the connection weight 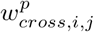 between neurons *i, j* in the different input SOMs are updated with a Hebbian learning rule as follows:

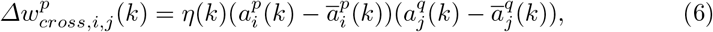

where

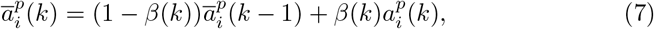

is a “momentum” like exponential decay and *η*(*k*), *β*(*k*) are monotonic (inverse time) decaying functions. Hebbian learning ensures that when neurons fire synchronously their connection strengths increase, whereas if their firing patterns are anti-correlated the weights decrease. The weight matrix encodes the co-activation patterns between the input layers (i.e. SOMs), as shown in Figure 3b, and, eventually, the learned growth law (i.e. relation) given the timeseries, as shown in Figure 5.

**Fig.5.**
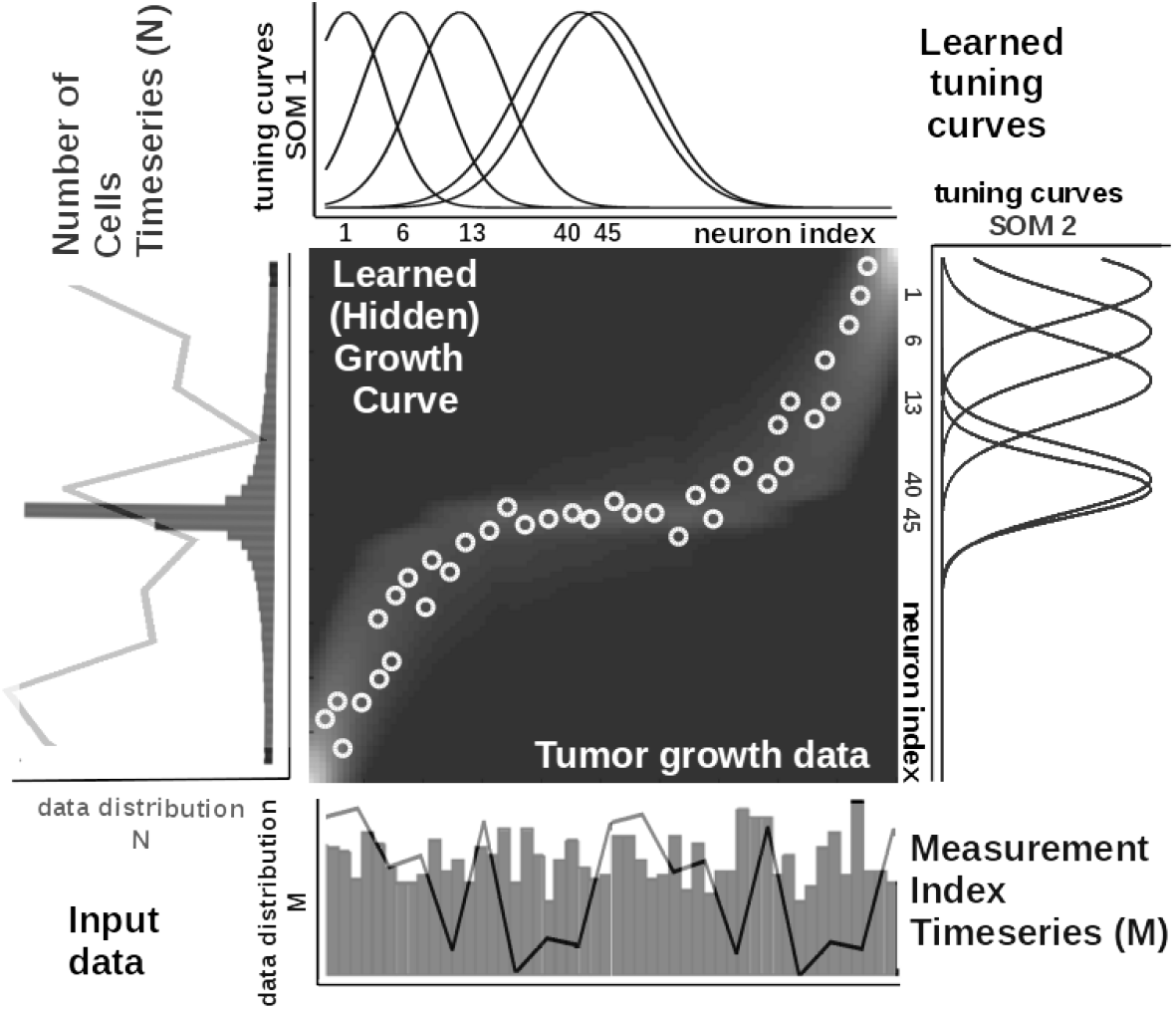
Extracted timeseries relation describing the growth law and data statistics for the data in Figure 3a depicting a cubic breast cancer tumor growth law among number of cells and irregular measurement over 150 weeks from [10]. Timeseries overlay on the data distribution and corresponding model encoding tuning curves shapes.

Self-organisation and Hebbian correlation learning processes evolve simultaneously, such that both the representation and the extracted relation are continuously refined, as new samples are presented. This can be observed in the encoding and decoding functions where the input activations are projected though *w_in_* (Equation 1) to the Hebbian matrix and then decoded through *w_cross_* (Equation 8).

#### Parametrization and read-out

In all of our experiments data from tumor growth timeseries is fed to the GLUECK network which encodes each timeseries in the SOMs and learns the underlying relation in the Hebbian matrix. The SOMs are responsible of bringing the timeseries in the same latent representation space where they can interact (i.e. through their internal correlation). In our experiments, each of the SOM has *N* = 100 neurons, the Hebbian connection matrix has size *NxN* and parametrization is done as: *alpha* = [0.01, 0.1] decaying, *η* = 0.9, 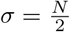 decaying following an inverse time law.

We use as decoding mechanism an optimisation method that recovers the real-world value given the self-calculated bounds of the input timeseries. The bounds are obtained as minimum and maximum of a cost function of the distance between the current preferred value of the winning neuron (i.e. the value in the input which is closest to the weight vector of the neuron in Euclidian distance) and the input sample at the SOM level. Depending on the position of the winning neuron in the SOM, the decoded / recovered value *y*(*t*) from the SOM neurons weights is computed as:

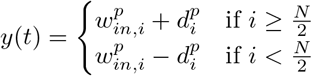

where, 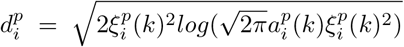 for the most active neuron with index *i* in the SOM, a preferred value 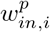 and 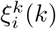 tuning curve size and 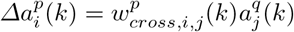. The activation 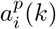 is computed by projecting one input through SOM *q* and subsequently through the Hebbian matrix to describe the paired activity (i.e. at the other SOM *p*, Equation 8).

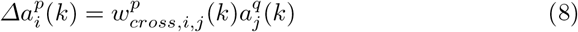

where 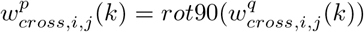 and *rot90* is a clockwise rotation.

### 2.2 Datasets

In our experiments we used publicly available tumor growth datasets (see Table 2), with real clinical tumor volume measurements, for breast cancer (datasets 1 and 2) and other cancers (e.g. lung, leukemia - datasets 3 and 4, respectively). This choice is to probe and demonstrate transfer capabilities of the models to tumor growth patterns induced by other cancer types.

**Table 2.** Description of the datasets used in the experiments.

| Experimental dataset setup |  |  |  |  |  |
| --- | --- | --- | --- | --- | --- |
| Dataset | Cancer Type | Data Type | Data Points | Data Freq. | Source |
| 1 | Breast (MDA-MB-231) | Fluorescence imaging | 7 | 2x/week | [20] |
| 2 | Breast (MDA-MB-435) | Digital Caliper | 14 | 2x/week | [26] |
| 3 | Lung | Caliper | 10 | 7x/week | [6] |
| 4 | Leukemia | Microscopy | 23 | 7x/week | [23] |

### 2.3 Procedures

In order to reproduce the experiments, the MATLAB ^®^ code and copies of all the datasets are available at https://gitlab.com/ ^1^ Each of the four mechanistic tumor growth models (i.e. Logistic, Bertalanffy, Gompertz, Holling) and GLUECK were presented the tumor growth data in each of the four datasets. When a dataset contained multiple trials, a random one was chosen.

#### Mechanistic models setup

Each of the four tumor growth models was implemented as ordinary differential equation (ODE) and integrated over the dataset length. We used a solver based on a modified Rosenbrock formula of order 2 that evaluates the Jacobian during each step of the integration. To provide initial values and the best parameters (i.e. *α, β*, λ, *k*) for each of the four models the Nelder-Mead simplex direct search (i.e. derivative-free minimum of unconstrained multi-variable functions) was used, with a termination tolerance of 10e^-6^ and upper bounded to 500 iterations. Finally, fitting was performed by minimizing the sum of squared residuals (SSR).

#### GLUECK setup

For GLUECK the data was normalized before training and de-normalized for the evaluation. The system was comprised of two input SOMs, each with *N* = 50 neurons, encoding the volume data and the irregular sampling time sequence, respectively. Both input density learning and cross-modal learning cycles were bound to 100 epochs. The full parametrization details of GLUECK are given in Parametrization and read-out section.

## 3 Results

In the current section we describe the experimental results, discuss the findings and also evaluate an instantiation of GLUECK learning capabilities to predict surgical tumor size.

All of the five models were evaluated through multiple metrics on each of the four datasets. This choice of different tumor types (i.e. breast, lung, leukemia) is to probe and demonstrate between and within tumor type prediction versatility. Assessing the distribution of the measurement error as a function of the caliper-measured volumes of subcutaneous tumors [5] suggested the following model for the standard deviation of the error *σ_i_* at each time point *i*,

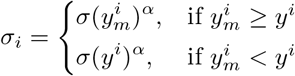

This formulation shows that, when overestimating (*y_m_ ≥ y*), the measurement error is *α* sub-proportional and, when underestimating (*y_m_ < y*), the error made is the same as the measured data points. In the following experiments we consider *α* = 0.84 and *σ* = 0.21, as a good trade-off of error penalty and enhancement. This interpretation of the notion of measurement error is used in the the metrics employed for prediction performance (i.e. Sum of Squared Errors (SSE), Root Mean Squared Error (RMSE), Symmetric Mean Absolute Percentage Error (sMAPE)) and goodness-of-fit and parsimony (i.e. Akaike Information Criterion (AIC) and Bayesian Information Criterion (BIC)), as shown in Table 3. Important to note that the evaluation metrics also take into account the number of parameters *p* each tumor growth model has, as in Table 1: Logistic *p* = 2, Bertalannfy *p* = 3, Gompertz *p* =2, Holling *p* = 3. GLUECK is a data-driven approach that does not use any biological assumptions or dynamical description of the process, rather it infers the underlying statistics of the data and uses it in predicting it without supervision. Note that the GLUECK hyper-parameters were unchanged during the experiments (i.e. the same network size, learning epochs etc.) and described in the Materials and methods. Due to its inherent learning capabilities GLUECK provides overall better accuracy between and within tumor type growth curve prediction. Assessing both summary statistics in Figure 6 and the broad evaluation in Table 4 we can conclude that a data-driven approach for tumor growth extraction can overcome the limitations that incomplete biological descriptions, the diversity of tumor types, and the small size of the data bring to mechanistic models.

**Table 3.**
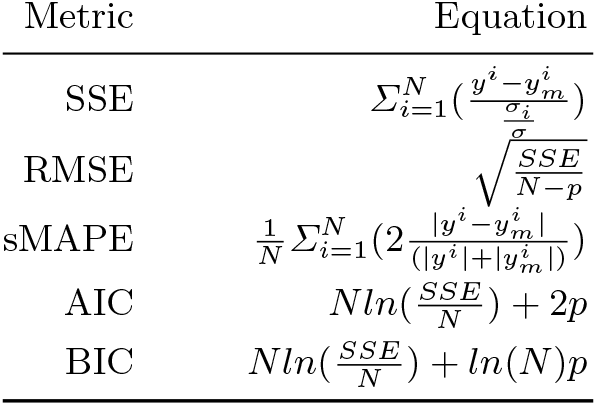
Evaluation metrics for tumor growth models. We consider: *N* - number of measurements, *σ* - standard deviation of data, *p* - number of parameters of the model.

**Fig.6.**
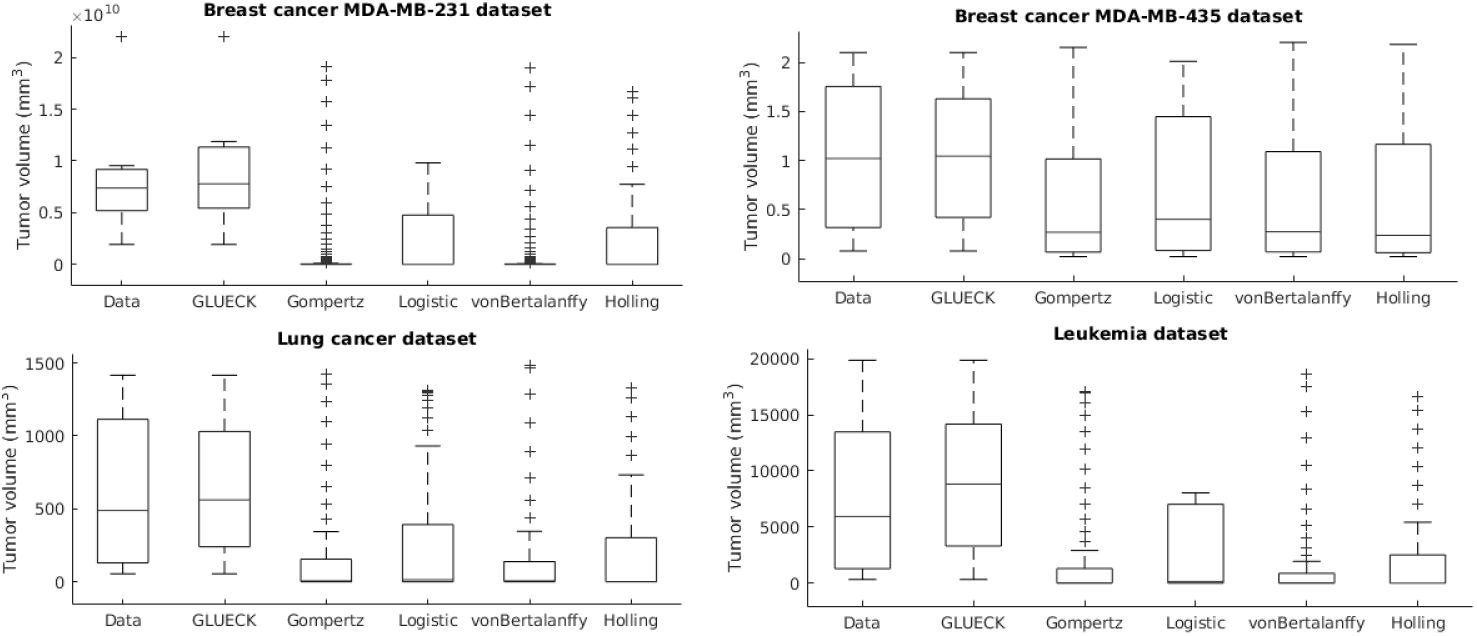
Evaluation of the tumor growth models: summary statistics.

**Table 4.** Evaluation of the tumor growth models.

Calculated as best in 3/5 metrics.<sup>b</sup> MDA-MB-231 cell line<sup>c</sup> MDA-MB-435 cell line
| Evaluation Metrics (smaller value is better) |  |  |  |  |  |  |
| --- | --- | --- | --- | --- | --- | --- |
| Dataset/Model | SSE | RMSE | sMAPE | AIC | BIC | Rank<sup>a</sup> |
| *Breast<sup>b</sup> cancer[20]* |  |  |  |  |  |  |
| Logistic | 7009.6 | 37.4423 | 1.7088 | 52.3639 | 52.2557 | **2** |
| Bertalanffy | 8004.9 | 44.7350 | 1.7088 | 55.2933 | 55.1310 | **5** |
| Gompertz | 7971.8 | 39.9294 | 1.7088 | 53.2643 | 53.1561 | **4** |
| Holling | 6639.1 | 40.7403 | 1.4855 | 53.9837 | 53.8215 | **3** |
| **GLUECK** | 119.3 | 4.1285 | 0.0768 | 19.8508 | 19.8508 | **1** |
| *Breast<sup>c</sup> cancer[26]* |  |  |  |  |  |  |
| Logistic | 0.2936 | 0.1713 | 0.1437 | -40.5269 | -39.5571 | **4** |
| Bertalanffy | 0.2315 | 0.1604 | 0.1437 | -41.3780 | -39.9233 | **2** |
| Gompertz | 0.3175 | 0.1782 | 0.1437 | -39.5853 | -38.6155 | **5** |
| Holling | 0.2699 | 0.1732 | 0.1512 | -39.5351 | -38.0804 | **3** |
| **GLUECK** | 0.0977 | 0.0902 | 0.0763 | -57.7261 | -57.7261 | **1** |
| *Lung cancer[6]* |  |  |  |  |  |  |
| Logistic | 44.5261 | 2.2243 | 1.5684 | 19.3800 | 20.1758 | **2** |
| Bertalanffy | 54.1147 | 2.6008 | 1.5684 | 23.5253 | 24.7190 | **5** |
| Gompertz | 53.2475 | 2.4324 | 1.5684 | 21.3476 | 22.1434 | **4** |
| Holling | 50.6671 | 2.5166 | 1.5361 | 22.8012 | 23.9949 | **3** |
| **GLUECK** | 3.6903 | 0.5792 | 0.2121 | -12.0140 | -12.0140 | **1** |
| *Leukemia[23]* |  |  |  |  |  |  |
| Logistic | 223.7271 | 3.2640 | 1.6368 | 56.3235 | 58.5944 | **2** |
| Bertalanffy | 273.6770 | 3.6992 | 1.6368 | 62.9585 | 66.3649 | **5** |
| Gompertz | 259.9277 | 3.5182 | 1.6368 | 59.7729 | 62.0439 | **4** |
| Holling | 248.5784 | 3.5255 | 1.6001 | 60.7461 | 64.1526 | **3** |
| **GLUECK** | 35.2541 | 1.2381 | 0.3232 | 9.8230 | 9.8230 | **1** |

## 4 Conclusion

Notwithstanding, tumor growth models have had a profound influence on modern chemotherapy, especially dose management, multi-drug strategies and adjuvant chemotherapy. Yet, the selection of the best model and its parametrization is not always straightforward. To tackle this challenge GLUECK comes as a support tool that could assist oncologists in extracting tumor growth curves from the data. Using unsupervised learning, GLUECK overcomes the limitations that incomplete biological descriptions, the diversity of tumor types, and the small size of the data bring on the basic, scalar growth models (e.g. Logistic, Bertalanffy, Gompertz, Holling). GLUECK exploits the temporal evolution of timeseries of growth data along with its distribution and obtains superior accuracy over basic models in extracting growth curves from different clinical tumor datasets. Without changes either to structure or parameters GLUECK’s versatility has been demonstrated in between cancer types (i.e. breast vs. lung vs. leukemia) and within cancer type (i.e. breast MDA-MB-231 and MDA-MB-435 cell lines) tumor growth curve extraction.

## Footnotes

1 The repository will be made visible after the review process.

